# Nutrient Abundance Signals the Changing of the Seasons by Phosphorylating PER2

**DOI:** 10.1101/2024.01.29.577255

**Authors:** Daniel C. Levine, Rasmus H. Reeh, Thomas McMahon, Thomas Mandrup-Poulsen, Ying-Hui Fu, Louis J. Ptáček

## Abstract

The circadian clock synchronizes metabolic and behavioral cycles with the rotation of the Earth by integrating environmental cues, such as light. Nutrient content also regulates the clock, though how and why this environmental signal affects the clock remains incompletely understood. Here, we elucidate a role for nutrient in regulating circadian alignment to seasonal photoperiods. High fat diet (HFD) promoted entrainment to a summer light cycle and inhibited entrainment to a winter light cycle by phosphorylating PER2 on serine 662. PER2-S662 phospho-mimetic mutant mice were incapable of entraining to a winter photoperiod, while PER2-S662 phospho-null mutant mice were incapable of entraining to a summer photoperiod, even in the presence of HFD. Multi-omic experimentation in conjunction with isocaloric hydrogenated-fat feeding, revealed a role for polyunsaturated fatty acids in nutrient-dependent seasonal entrainment. Altogether, we identify the mechanism whereby nutrient content shifts circadian rhythms to anticipate seasonal photoperiods in which that nutrient state predominates.

**HIGHLIGHTS:**

- High fat diet promotes entrainment to summer but inhibits entrainment to winter.
- Calorie restriction promotes entrainment to winter but inhibits entrainment to summer.
- PER2-S662 phosphorylation is required for nutritional regulation of seasonal circadian entrainment.
- Dietary polyunsaturated fatty acids regulate seasonal circadian entrainment.

## INTRODUCTION

Circadian clocks have evolved to anticipate 24-hour cycles of light and dark that arise from rotation of the Earth. A key feature of the clock is that periodicity is not exactly 24 hours and thus requires constant re-synchronization with the external environment from periodic and repetitive environmental cues, such as light. A surprising finding has been that nutrient content also serves as a clock-synchronizing cue for mammals ^1,2^, though nutrient availability is not necessarily periodic or repetitive. A major goal of ongoing research is to interrogate the mechanisms and contexts that integrate the molecular clock with external timing cues, especially nutrition.

The rotation of the Earth is tilted relative to the orbital plane, which causes yearly patterns in the intensity and daily duration of light that, in turn, drives cycles of nutrient abundance and limitation. Decreased light duration and solar energy during the winter is associated with decreased plant biomass and adaptations such as hibernation and migration in animals. Conversely, longer light duration and more intense solar energy during the summer is associated with plant growth and adaptations such as gorging and reproductive behavior in animals ^3,4^. In a laboratory setting, mice exposed to seasonal light cycles demonstrate profound reprogramming of peripheral tissue gene expression and feeding behavior ^5^. Humans display seasonal variations in daily activity, body weight, metabolic biomarkers, and genome-wide transcription, though societal and technological factors likely confound some seasonal phenotypes ^6–9^.

Most research is performed with a model day consisting of alternating cycles of 12 hours of light and 12 hours of dark (12:12 LD). In nature, however, the photoperiod is not constant and organisms must continually phase-shift internal rhythms to maintain synchrony with the external environment. Nocturnal organisms, such as mice, must phase-advance each day until the winter solstice occurs and phase-delay each day until the summer solstice occurs. While the neurons that regulate circadian rhythms in *Drosophila melanogaster* are essential for aligning behavioral rhythms with seasonal light cycles ^10^, a gap remains in our understanding of the mechanisms whereby the mammalian clock adapts to seasonal light cycles and of the role that coincident changes in nutrient availability play in that process.

Genetic analysis of humans with familial advanced sleep phase (FASP), whose daily activity is shifted relative to the prevailing light/dark cycle, uncovered a phosphorylation event on the circadian repressor, PER2 ^11^. Mice and humans harboring a phospho-null mutation wherein serine 662 is mutated to a glycine (*PER2-S662G*) have advanced behavioral phase and a shortened circadian period, while mutation to aspartate (*PER2-S662D*) that mimics the phosphorylated state does the opposite ^12^. PER2-S662 is phosphorylated by casein kinase (CK1δ) but integrates numerous nutrient-signaling pathways that either modify PER2-S662 with alternative marks or drive competitive modifications at nearby sites ^13–16^. Here, we interrogate the extent to which nutrient shifts circadian rhythms to anticipate changing seasonal photoperiods by phosphorylating PER2-S662.

We found that mice fed a high fat diet were defective in advancing daily behavioral rhythms to align with the longer night-length of winter, but more readily delayed daily behavioral rhythms to align with the shorter night-length of summer. Mice on a calorie restricted diet had the opposite phenotype, suggesting that caloric content or nutrient composition of chow drives entrainment to the season in which that nutrient state predominates. PER2-S662 phosphorylation in the hypothalamus was increased by high fat feeding, while mice harboring a *PER2-S662G* phospho-null mutation rapidly entrained to a winter photoperiod even when fed a high fat diet. Mechanistically, acute hypo-nutritive challenge decreases PER2-S662 phosphorylation in the hypothalamus to transcriptionally reprogram polyunsaturated fatty acid metabolism. Partially hydrogenating the fat source in mouse chow to convert polyunsaturated fatty acids into monounsaturated fatty acids promoted circadian entrainment to a summer light cycle and inhibited entrainment to a winter light cycle, suggesting that these essential fatty acids play a key role in circadian adaptation to nutrient availability. Altogether, we reveal PER2-S662 phosphorylation and transcriptional control of polyunsaturated fatty acid metabolism as a key mechanism whereby mice maintain synchrony between the nutritive state, the external light/dark cycle, and circadian rhythms across the year.

## RESULTS

### Nutrient density controls circadian entrainment to a winter light cycle

Interventions that cause nutrient excess or limitation in mice regulate clock function by altering free-running behavioral period, the diurnal distribution of wheel-running activity, and the peripheral circadian transcriptome ^1,2,17,18^. As yearly cycles in nutrient availability are coincident with the daily duration of light on much of the Earth, we sought to interrogate the extent to which nutrient density regulates entrainment of circadian rhythms to seasonal light cycles.

During the winter, the daily dark period is longer and nutrient content becomes limited across most of the planet. We sought to test the extent to which nutrient composition regulates the ability of mice, a nocturnal species, to advance daily behavioral rhythms to entrain to a winter light cycle. We first gave wild type mice *ad libitum* access to either a calorie-dense, high fat diet (“HFD”, 4.6 kcal/g, 45% calories from fat) or regular chow (“Reg”, 3 kcal/g, 13% calories from fat) and monitored *ad libitum* wheel-running activity after shifting lights from alternating cycles of 12 hours of light and 12 hours of dark (12:12 LD), which approximates the equinox, to alternating cycles of 4 hours of light and 20 hours of dark (4:20 LD), which approximates winter (**Figure 1A**). We chose to simulate a winter light cycle by asymmetrically advancing the dark period to conserve “*Zeitgeber-time*” (ZT) field-standard nomenclature that is relative to the onset of the light period. In a 12:12 LD cycle, the onset of daily behavioral rhythms in wild type mice occurs at the time when the lights turn off, ZT12. After shifting lights to 4:20 LD, wild type mice advanced daily behavioral rhythms towards the new lights-off time (ZT4) at a rate of ∼0.25 hours/day (**Figure 1B and Supplemental Figure 1A**). Strikingly, mice given *ad libitum* access to HFD had a slower rate of behavioral entrainment to the 4:20 LD photoperiod that was equivalent to approximately half the rate of mice on regular chow (**Figure 1B and Supplemental Figure 1A**). Indeed, 30 days after shifting the light cycle to approximate winter, mice on HFD had accumulated a 2-hour phase-delay in activity onset relative to mice on regular chow (**Supplemental Figure 1B**). Thus, HFD decreases the rate at which mice entrain to a winter light cycle.

**Figure 1:**
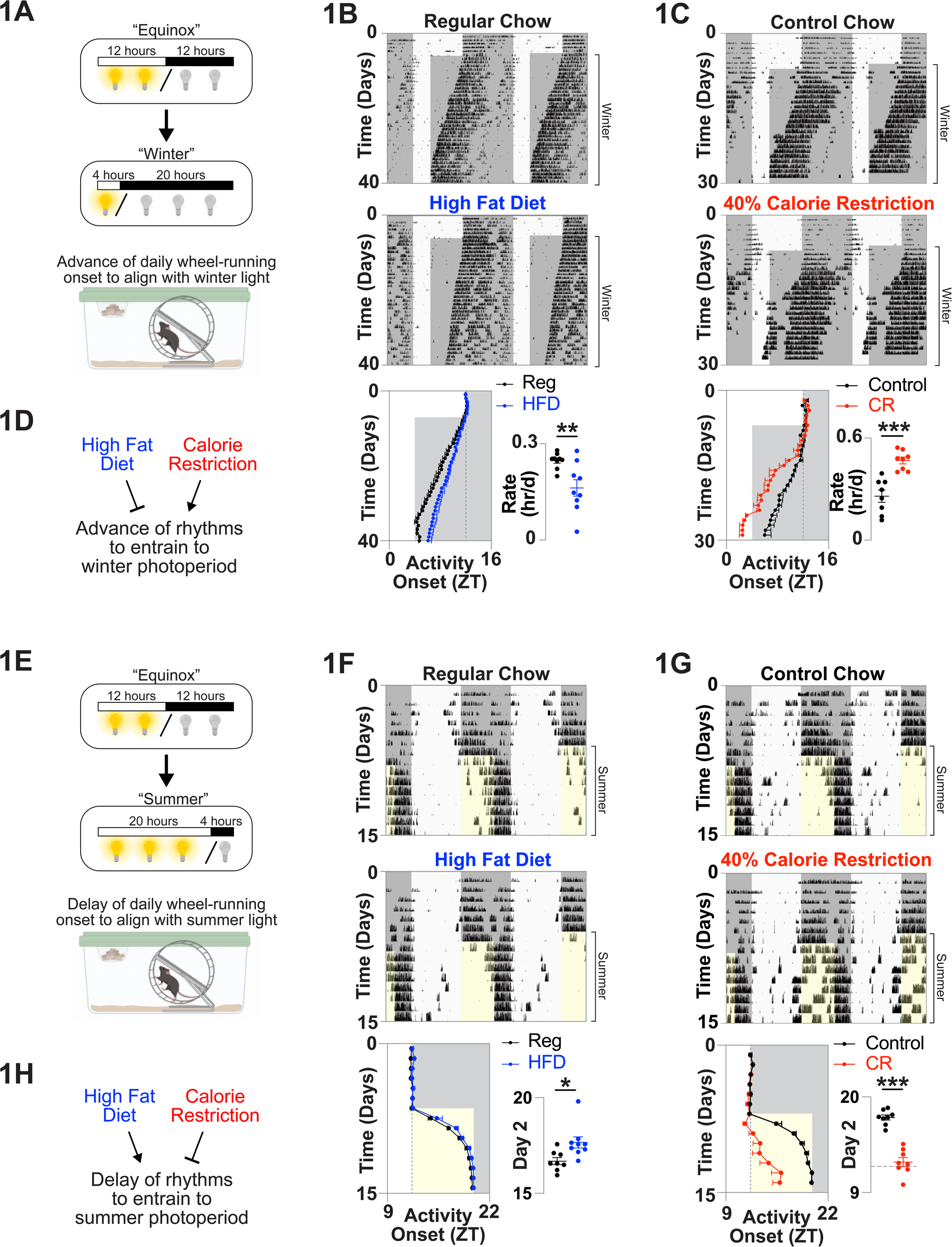
Nutrient density controls seasonal circadian entrainment. **A)** Model of winter entrainment assay. **B and C)** Representative, double-plotted wheel-running traces from mice shifted from a 12:12 LD cycle to a 4:20 LD cycle and given **(B)** *ad libitum* regular chow or high fat diet or **(C)** control chow or a calorie restricted diet. Quantifications of average activity onset and rate of behavioral advance for all replicates. **D)** Model of results. **E)** Model of summer entrainment assay. **F and G)** Representative, double-plotted wheel-running traces from mice shifted from a 12:12 LD cycle to a 20:4 LD cycle and given **(F)** *ad libitum* regular chow or high fat diet or **(G)** control chow or a calorie restricted diet. Quantifications of average activity onset across the duration of the recording and activity onset on the 2^nd^ day after shifting the lights for all replicates. **H)** Model of results. *p<0.05, **p<0.01, ***p<0.001. Data are represented as mean +/-SEM. See also Supplemental Figure 1.

We next interrogated whether calorie deficiency altered the rate at which mice entrain to a winter light cycle. We utilized computer-assisted feeder devices to give mice either 16, 300-mg pellets of a control chow (BioServ, AIN-93M) or 10, 300-mg pellets of a macronutrient-controlled diet (BioServ, 40% DR AIN-93M) to achieve a 40% calorie restriction (CR). We controlled for elongated fasting durations that occur during traditional CR paradigms ^2^ by programming the computer to dispense food at even intervals from ZT4 to ZT24 (every 2 hours for CR mice and every 1.25 hours for control mice) (**Supplemental Figure 1C**). Under this paradigm, wild type mice on control chow advanced their daily behavioral rhythms to align with the 4:20 LD cycle that approximates winter at a rate of ∼0.25 hours/day, while mice on CR entrained faster, at a rate that was nearly double that of the mice on control chow (**Figure 1C and Supplemental Figure 1D**). Twenty days after shifting the lights to 4:20 LD, mice on CR were phase-advanced from mice on the control chow by 4 hours (**Supplemental Figure 1E**). Thus, CR increases the rate at which mice entrain to a winter light cycle. These data show that HFD and CR have opposite effects on the rate at which mice advance daily behavioral rhythms to align with the winter photoperiod (**Figure 1D**).

### Nutrient density controls circadian entrainment to a summer light cycle

During the summer, the daily dark period is shorter and nutrients are more abundant across much of the planet. We next tested whether nutrient density affects the ability of mice to delay daily behavioral rhythms to entrain to a summer light cycle. We gave mice HFD or regular chow and monitored wheel-running activity after shifting lights from 12:12 LD to alternating cycles of 20 hours of light and 4 hours of dark (20:4 LD), which approximates summer (**Figure 1E**). Again, we chose to asymmetrically shift light cycles to maintain ZT references that are standard in the field. We note that the kinetics of entrainment are faster for photoperiods with extended light duration due, in part, to behavioral masking of nocturnal animals by light ^19^. Two days after shifting to a summer light cycle, mice on regular chow had delayed behavioral onset by 4.8 hours, equivalent to 60% of the shift that is required to align with the new lights-off time (ZT20). By 4 days after shifting the light cycle, mice on regular chow were entrained to within one hour of the new lights-off time (ZT20). Examining the effect of HFD revealed a significantly faster behavioral entrainment to a summer light cycle relative to mice on regular chow (**Figure 1F and Supplemental Figure 1F**). Indeed, 2 days after changing the light cycle to 20:4 LD, the activity onset of HFD mice was delayed by 5.8 hours, which is one hour more than mice on regular chow or 12.5% more entrained to the new lights-off time (**Figure 1F**). These results reveal that HFD promotes entrainment to a summer light cycle, which is in striking contrast to our observation that HFD inhibits entrainment to a winter light cycle (**Figure 1B**).

We next interrogated the capacity of nutrient deficiency to alter the rate at which mice entrain to a summer light cycle. Here, we administered control and CR chows at even intervals throughout the entire day (from ZT0-ZT24) to ensure similar feeding/fasting patterns for both groups of mice (**Supplemental Figure 1G**). Interestingly, CR slowed the rate at which mice entrained to the 20:4 LD cycle that approximates summer (**Figure 1G and Supplemental Figure 1H**), which is in contrast to our observation that CR promotes entrainment to a winter light cycle (**Figure 1C**). Mice on the CR diet delayed behavioral onset by only 0.5 hours within 2 days and only 3.65 hours by the end of the recording window, equivalent to less than 50% of the required shift (**Figure 1G**). In contrast, mice on the control diet had delayed behavioral onset by 5.6 hours after 2 days and were fully entrained at the end of the recording window (**Figure 1G**). These experiments reveal that CR decreases the rate at which mice phase-delay behavioral rhythms to align to a summer light cycle while HFD increases it (**Figure 1H**). These experiments suggest that hyper-and hypo-caloric conditions, and/or associated changes in nutrient composition and organismal nutrient stores, drive circadian entrainment to the season in which those nutrient conditions predominate and inhibit seasonal entrainment if nutrient is unseasonably available.

### PER2 dosage regulates seasonal circadian entrainment

While clock neurons in *Drosophila melanogaster* regulate seasonal circadian adaptation ^10^, to our knowledge, no genetic studies have been performed in mammals to specifically interrogate the role of the circadian clock in entrainment to seasonal light cycles. To understand the extent to which the genetic clock machinery affects the rate at which mice entrain to seasonal light cycles, we assayed two genetic mutant mouse lines: the first in which the mouse core clock gene, *Per2*, was deleted (*Per2 ^-/-^*) ^20^; and the second in which five copies of the human *PER2* gene fused to a C-terminal FLAG tag are stably integrated into the genome under endogenous proximal regulatory elements using BAC transgenesis (*PER2-TgWT*). *Per2* knockout, *PER2-TgWT*, and wild type littermates were housed in wheel-cages and daily wheel-running behavior was monitored before and after shifting lights to a 4:20 LD cycle that approximates winter. Wild type mice advanced daily activity onset to align with the winter light cycle in 35 days. Interestingly, PER2 dosage affected the rate at which mice entrained to a winter light cycle, as *PER2-TgWT* mice failed to shift activity patterns within 60 days of monitoring wheel running whereas *Per2* knockout mice entrained quickly, in an average of 13 days (**Figure 2A**). Indeed, *Per2* knockout mice entrained at a rate of 0.48 hours/day while *PER2-TgWT* mice entrained at a rate of 0 hours/day (**Figure 2A**). 30 days after shifting the light cycle to approximate winter, *Per2* knockout mice demonstrated an ∼7.5 hour phase-advance relative to the *PER2-TgWT* mice in average (**Supplemental Figure 2A**).

**Figure 2:**
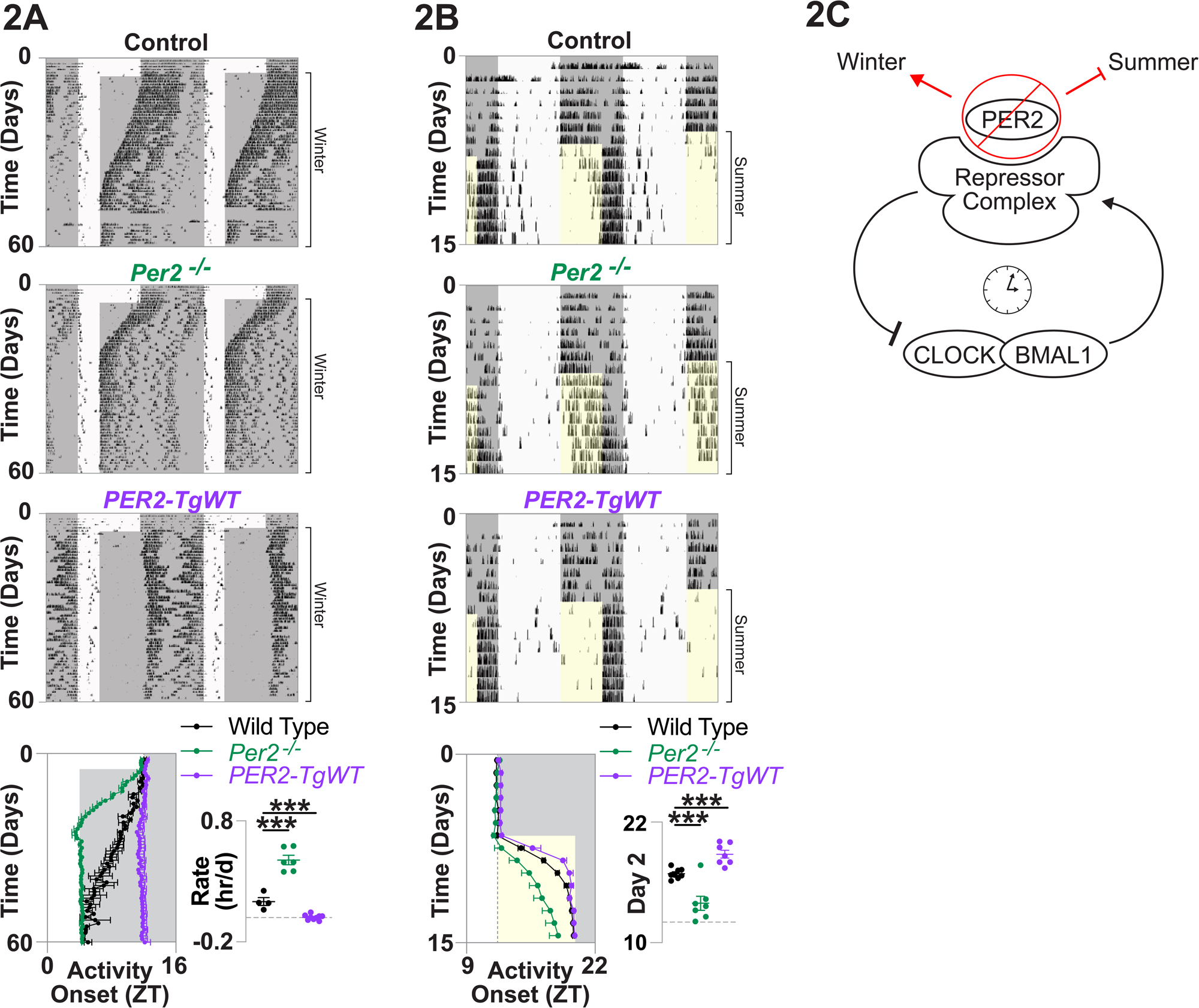
PER2 dosage regulates seasonal circadian entrainment. **A and B)** Representative, double-plotted wheel-running traces from *Per2* knockout, *PER2-TgWT* overexpressing, and wild type littermate controls that were shifted from a 12:12 LD cycle to **(A)** a 4:20 LD cycle that approximates winter or **(B)** a 20:4 LD cycle that approximates summer. Bottom panels: quantification of average activity onset for each day of the recording and **(A)** rates of behavioral advance and **(B)** activity onset on the 2^nd^ day for each replicate. **C)** Model of results depicting effect of *Per2* knockout on seasonal entrainment. ***p<0.001. Data are represented as mean +/-SEM. See also Supplemental Figure 2.

We next assessed the extent to which PER2 dosage regulated entrainment to a summer light cycle by assaying wheel-running activity of the mutant mice before and after shifting the light cycle to 20:4 LD. Here, *PER2-TgWT* mice entrained faster to the summer light cycle than wild type controls, while *Per2* knockout mice entrained slower (**Figure 2B**). Two days after changing the light cycle, *PER2-TgWT* mice had delayed activity onset by 6.8 hours to ZT18.8, 2 hours more than wild type controls (**Figure 2B**). In contrast, *Per2* knockout mice had delayed activity onset by just 1.9 hours within 2 days and failed to delay activity onset beyond 4.6 hours over the 15-day window where we monitored wheel-running activity (**Figure 2B**). Together, these data show that genetically regulating PER2 dosage controls the rate at which mice phase-shift behavioral rhythms to align with seasonal light cycles such that low PER2 levels promote behavioral entrainment to winter and inhibit behavioral entrainment to summer (**Figure 2C**).

### HFD increases PER2-S662 phosphorylation to regulate seasonal circadian entrainment

We previously identified a phosphorylation site on serine 662 of PER2 that is mutated to a glycine (*PER2-S662G*) in humans with familial advanced sleep phase (FASP) ^11^. *PER2-S662G* abrogates a casein kinase 1δ (CK1δ)-regulated phosphorylation site on PER2 and reduces PER2 protein level. Mutation of PER2-S662 to aspartate (*PER2-S662D*) to mimic the phosphorylated state delays sleep phase in mice and increases protein level, suggesting that this phosphorylation event is sufficient for regulating behavioral phase in mice and humans ^12^. PER2 post-translational modifications that compete with PER2-S662 phosphorylation have been identified from nutrient signaling pathways including O-GlcNAc transferase (OGT), glycogen synthase kinase (GSK3β), and the mammalian homolog of the silent information regulator two (SIRT1) ^13–16^. As PER2 dosage regulates entrainment to seasonal light cycles, we hypothesized that PER2-S662 phosphorylation constitutes an epigenetic mechanism to alter circadian phase in response to seasonal light cycles and nutrient availability.

We first sought to determine the extent to which PER2-S662 phosphorylation alters the rate at which mice entrain to seasonal light cycles. Here, we utilized mouse lines that express human *PER2* under endogenous proximal regulatory elements but wherein S662 is mutated to either a glycine (*PER2-S662G*) to ablate the phosphorylation site, or to an aspartate (*PER2-S662D*) to mimic the phosphorylated state (**Figure 3A**) ^12^. We monitored wheel-running activity from *PER2-S662G*, *PER2-S662D*, and wild type littermates before and after the lights were shifted to a 4:20 LD light cycle that approximates winter. While wild type mice entrained to a 4:20 LD light cycle in ∼35 days, *PER2-S662D* phospho-mimetic mice failed to shift the onset of daily wheel-running behavior within the 60 days that we monitored wheel-running activity (**Figure 3B**). In contrast, *PER2-S662G* phospho-null mice entrained much more quickly, advancing their daily activity onset to align with the new lights-off time within just 4 days (**Figure 3B**). Indeed, *PER2-S662G* mice advanced daily activity onset at an average rate of ∼1.25 hours/day, while *PER2-S662D* mice advanced at 0 hours/day (**Figure 3B**). Thirty days after shifting the light cycle to approximate winter, *PER2-S662G* mice had accumulated a phase-advance of ∼8 hours relative to *PER2-S662D* mice (**Supplemental Figure S3A**).

**Figure 3:**
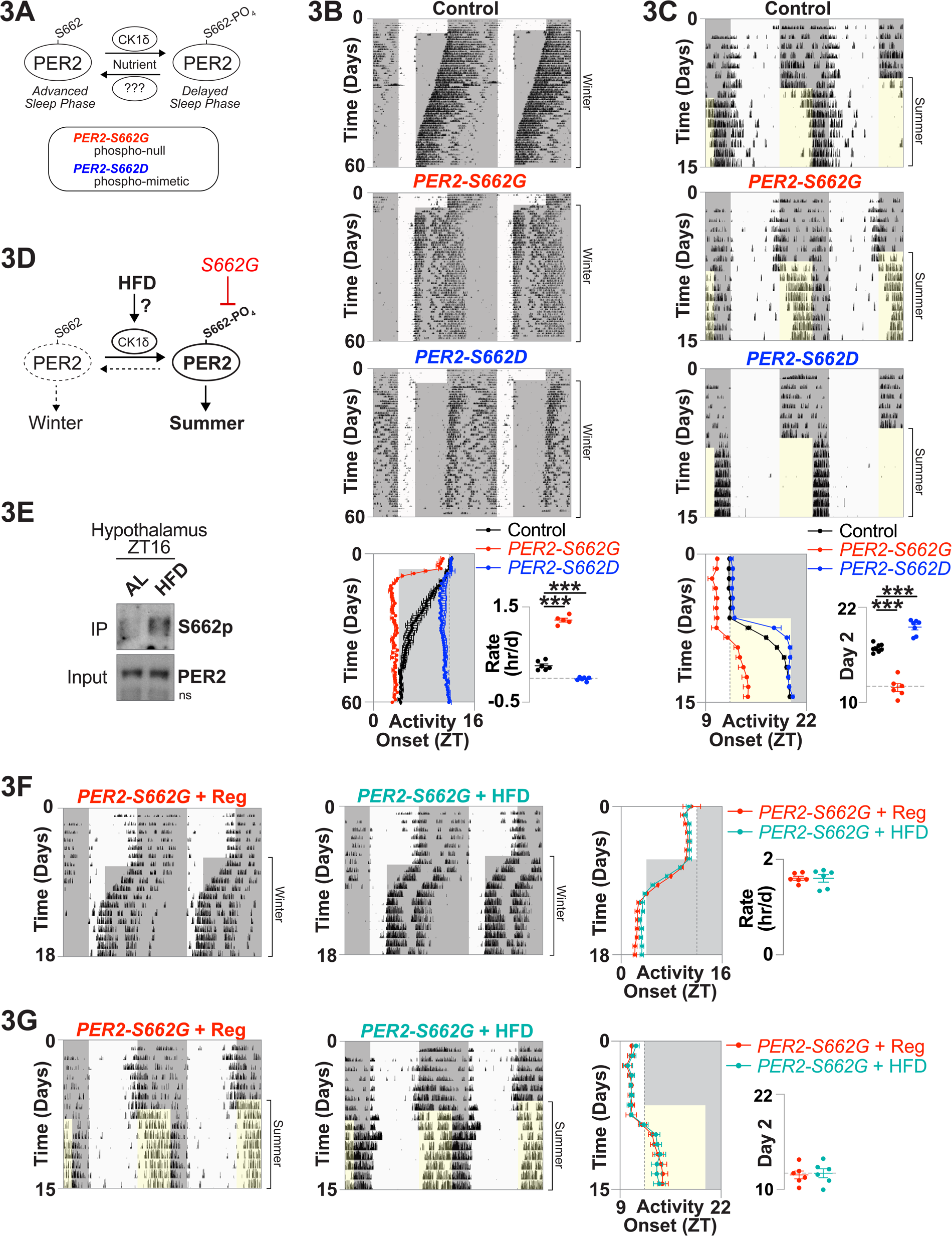
HFD increases PER2-S662 phosphorylation to regulate seasonal circadian entrainment. **A)** Model depicting genetic and biochemical regulation of PER2-S662 phosphorylation. **B and C**) Representative wheel-running traces and quantifications for **(B)** winter entrainment and **(C)** summer entrainment assays for *PER2-S662G*, *PER2-S662D*, and wild type littermate controls. **D)** Experimental model hypothesizing convergent regulation of seasonal entrainment by HFD and PER2-S662 hyper-phosphorylation. **E)** Immunoprecipitation of FLAG-tagged PER2 and immunoblotting for PER2-S662 phosphorylation and total PER2 from hypothalamus of *PER2-TgWT* mice collected at ZT16 following 1 week of HFD feeding. **F and G)** Representative, double-plotted wheel-running traces from *PER2-S662G* mice given *ad libitum* access to regular chow or HFD and that were shifted from 12:12 LD to **(F)** 4:20 LD or **(G)** 20:4 LD and quantified as previously. ***p<0.001. ns = non-specific band. Data are represented as mean +/-SEM. See also Supplemental Figure 3.

We next determined the effect of PER2-S662 phospho-mutants on the ability of mice to entrain to a 20:4 LD light cycle that approximates summer. Similar to our previous observations with nutrient challenge (**Figure 1**) and genetic interrogation of PER2 dosage (**Figure 2**), PER2-S662 phospho-mutants had the opposite effect on entrainment to a summer light cycle than they did on a winter light cycle. *PER2-S662D* mice entrained quickly to the summer light cycle while *PER2-S662G* failed to entrain within the experimental window (**Figure 3C**). Indeed, 2 days after shifting the light cycle to approximate summer, *PER2-S662D* phospho-mimetic mice demonstrated an average activity onset of ZT19.5, constituting >90% of the required entrainment within this short time window (**Figure 3C**). In contrast, *PER2-S662G* phospho-null mice demonstrated an activity onset of just ZT12 two days after shifting the light cycle and failed to delay beyond ZT14.5 over the duration of the recording (**Figure 3C**). Together, these results indicate that PER2-S662 phosphorylation regulates behavioral entrainment to seasonal light cycles such that *PER2-S662G* drives phase-shifts that promote entrainment to a winter light cycle but inhibit entrainment to summer.

HFD decreases the rate at which mice entrain to a winter light cycle and increases the rate at which they entrain to a summer light cycle (**Figure 1**). As PER2-S662 phosphorylation is influenced by nutrient-signaling pathways ^13–16^ and pharmacologic inhibition of CK1δ during HFD feeding improves circadian rhythms in 12:12 LD ^21^, we hypothesized that HFD affects seasonal entrainment by increasing PER2-S662 phosphorylation (**Figure 3D**). We first assessed hypothalamic PER2-S662 phosphorylation at ZT16, the peak in PER2, from *PER2-TgWT* mice housed in 12:12 LD cycles and given *ad libitum* access to HFD for one week. Of note, hypothalamic dissections were made with a brain matrix using the coordinates (relative to bregma): 0 to -3 mm anteroposterior, 1.5 to -1.5 mm mediolateral, 4 to 7 mm dorsoventral. Thus, lysates contain the SCN, the paraventricular nucleus, the arcuate nucleus, and other hypothalamic nuclei that are relevant for basal metabolic behaviors. PER2 was immunoprecipitated from the hypothalamic lysates with anti-FLAG antibodies, and PER2-S662 phosphorylation was monitored with site-specific antibodies by Western blotting ^14^. We observed increased PER2-S662 phosphorylation in the hypothalamus following 1 week of HFD feeding (**Figure 3E**), in line with previous studies suggesting increased CK1δ activity during HFD feeding ^21^. Increased PER2-S662 phosphorylation by HFD is also consistent with our observations that *PER2-S662D* and HFD both decrease the rate at which mice entrain to a winter photoperiod and increase the rate of entrainment to a summer photoperiod (**Figures 1B, 1F, and 3B-C**).

We next sought to interrogate whether increased PER2-S662 phosphorylation during HFD was necessary for altering circadian entrainment to seasonal light cycles. To test this, phospho-null, *PER2-S662G* mice were given *ad libitum* access to regular chow or a HFD and shifted to either a 4:20 LD cycle that approximates winter, or to a 20:4 LD cycle that approximates summer (**Figure 3D**). We had previously seen that HFD halved the rate at which wild type mice entrain to a winter light cycle and increased the rate at which wild type mice entrained to a summer light cycle (**Figures 1B and 1F**). Intriguingly, HFD had no effect on the rate at which *PER2-S662G* mice entrained to summer or winter light cycles, suggesting that HFD regulates seasonal circadian entrainment by increasing phosphorylation of PER2-S662. Specifically, we observed that *PER2-S662G* mice entrained to a 4:20 LD cycle that approximates winter within 4 days, regardless of whether they had access to regular chow or HFD (**Figure 3F**). Similarly, HFD did not affect activity onset profiles of *PER2-S662G* mice after shifting lights to a 20:4 LD cycle that approximates summer as neither dietary condition had phase-delayed beyond ZT14.5 within the recording window and were thus ∼5.5 hours phase-advanced from lights-off time (**Figure 3G**). Altogether, these experiments reveal that HFD regulates circadian entrainment to seasonal photoperiods by increasing PER2-S662 phosphorylation.

### Fasting decreases hypothalamic PER2-S662 phosphorylation to regulate behavior

We next sought to interrogate the mechanism whereby PER2-S662 phosphorylation affects hypothalamic clock function during nutrient challenge. We turned to an acute fasting paradigm in standard, 12:12 LD cycles to isolate molecular effects from differences in circadian phase that are inherent to PER2 mutant animals during seasonal entrainment. During fasting, mice enter a transient period of inactivity and energy conservation termed ‘torpor’ that is regulated by hypothalamic nuclei ^22,23^ and the circadian clock ^24^. Signals that activate CK1δ, such as insulin and gastrin, are low during fasting ^25,26^, suggesting that PER2-S662 is hypo-phosphorylated in this condition.

We sought to interrogate the extent to which decreased levels of PER2-S662 phosphorylation drive the molecular and behavioral response to fasting (**Figure 4A**). We first measured PER2-S662 phosphorylation in the hypothalamus after 16 hours of fasting, the time where PER2 protein reaches its maximum and torpor is beginning ^24^. Immunoprecipitation of hypothalamic FLAG-tagged PER2 from *PER2-TgWT* mice and blotting for PER2-S662 phosphorylation revealed decreased PER2-S662 phosphorylation during fasting (**Figure 4B**). Next, we tested the extent to which PER2-S662 phosphorylation, and the clock more generally, play a role in the behavioral response to fasting. We monitored *ad libitum* fed and fasted activity in *PER2-S662D* phospho-mimetic mutant, *Per2* knockout, and control mice housed in a video-based activity tracking apparatus. During *ad libitum* feeding, *PER2-S662D* had similar activity to control mice during the late-dark period (yellow bar), while *Per2* knockout mice had a non-significant trend towards decreased activity (**Figures 4C and 4E and Supplemental Figure 4A**). While fasting decreased activity in wild type mice within the late-dark period, *PER2-S662D* mice had markedly increased activity (18-24 hours of fasting, orange bar) that repeated at the same circadian phase in the 2^nd^ day of fasting (42-48 hours of fasting, red bar) (**Figures 4D-E**). Within the same time windows, fasting decreased activity in *Per2* knockout mice relative to wild type littermates (**Supplemental Figures 4B-C**). These results show that fasting decreases PER2-S662 phosphorylation and that this event is necessary for down-regulating behavior at the time when torpor is induced ^24^.

**Figure 4:**
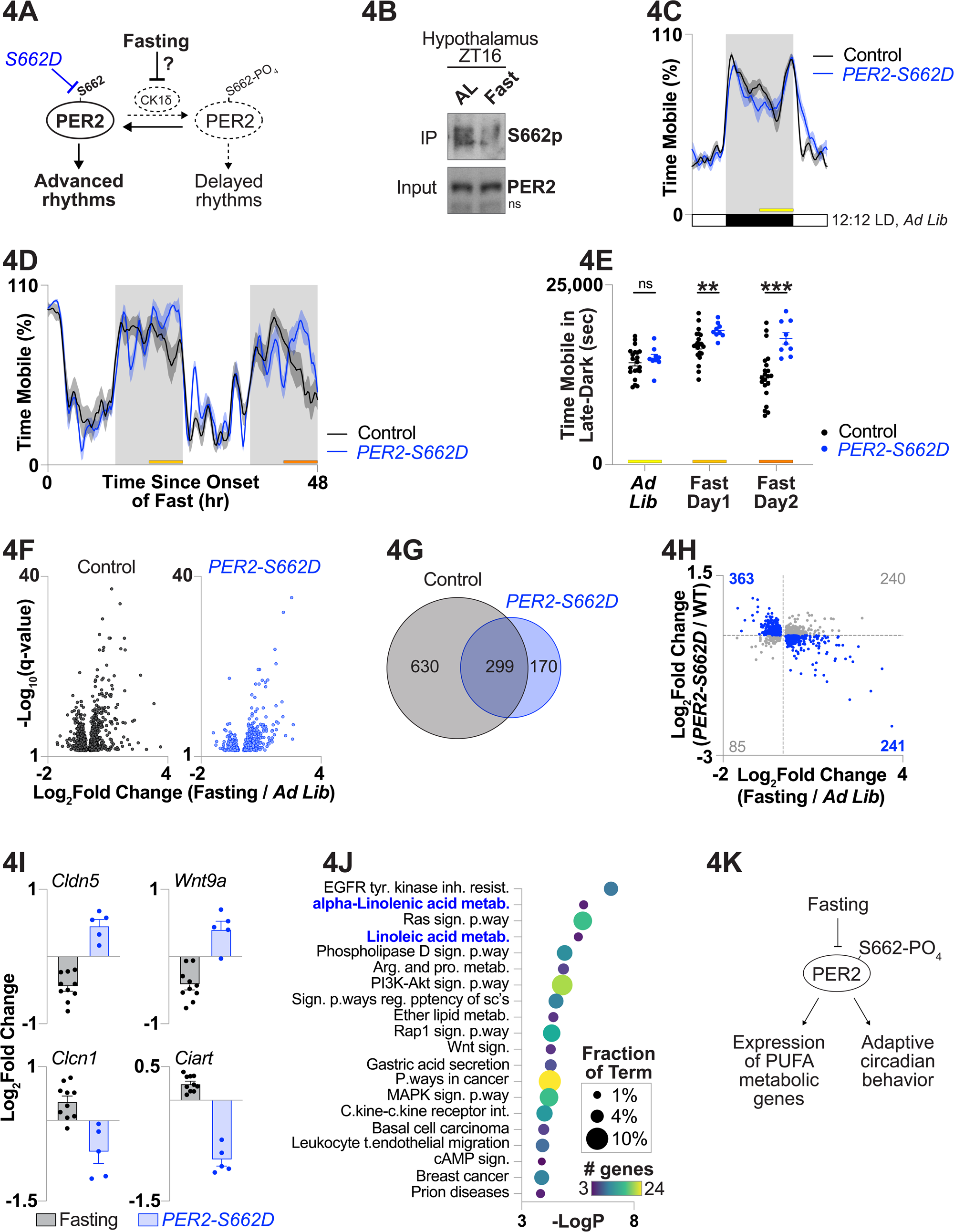
Fasting decreases PER2-S662 phosphorylation to regulate hypothalamic transcription and behavior. **A)** Experimental model hypothesizing convergent regulation of circadian behavior by fasting and PER2-S662 hypo-phosphorylation. **B)** Immunoprecipitation of FLAG-tagged PER2 and immunoblotting for PER2-S662 phosphorylation and total PER2 from hypothalamus of *PER2-TgWT* mice collected at ZT16 following 16 hours of fasting. **C)** Average time that mice were mobile within each 15 minute window across a standard 12:12 LD cycle for *ad libitum* fed *PER2-S662D* mice and wild type littermate controls using video-based activity tracking. Bar indicates ZT18-24**. D)** Average mobile time for *PER2-S662D* mice and wild type littermate controls where food was removed from the cage at ZT0. Bars indicate ZT18-24 in the first and second days of fasting. **E)** Quantification of late-dark (ZT18-24) mobile time in *ad libitum* fed and fasted *PER2-S662D* and control mice. **F)** RNA-sequencing from fed and fasted, control and *PER2-S662D* hypothalamus collected at ZT16. Volcano plots depicting differentially expressed genes (DESeq2 FDR-adjusted p-value<0.05) by fasting in each genotype and **(G)** overlap. **H)** Quadrant plot comparing the transcriptional response to fasting in wild type mice (x-axis) and the transcriptional response to *PER2-S662D* in fasted mice (y-axis) for hypothalamic genes that are differentially expressed by fasting in wild type mice at ZT16. Genes within quadrants 2 and 4 are highlighted in blue with **(I)** examples shown. **J)** gene ontology analysis of top-enriching KEGG terms for genes within quadrants 2 and 4 from panel H above. Terms associated with polyunsaturated fatty acid metabolism are highlighted in blue. **K)** Model of results. **p<0.01, ***p<0.001. Data are represented as mean +/-SEM. See also Supplemental Figure 4 and Supplemental Table 1.

### *PER2-S662D* opposes the transcriptional response to fasting in the hypothalamus

PER2 is part of a megadalton circadian repressor complex that, together with other core-clock and collaborative transcription factors, drive rhythmic expression of ∼10% of the transcriptome in any given cell ^27–29^. The oscillatory transcriptome is context-specific and has been shown to be reprogrammed in peripheral tissues by diverse nutrient challenges, including fasting ^30^. Thus, we next sought to characterize the hypothalamic transcriptome that is regulated by PER2-S662 phosphorylation during fasting. We excised hypothalamus from *ad libitum* fed and 16-hour fasted wild type and *PER2-S662D* mice at ZT16, the peak in PER2, and conducted RNA-sequencing. We found that 929 genes were differentially expressed (“DE”, DESeq2 FDR-adjusted p-value <0.05) by fasting compared with *ad lib* fed in wild type mice, but only 469 genes were DE by fasting compared with *ad lib* fed in the *PER2-S662D* mice, of which 299 DE genes were shared across both genotypes (**Figures 4F-G and Supplemental Table 1**). We also observed an inverse relationship between the transcriptional response to fasting in wild type mice and the transcriptional response to the *PER2-S662D* genotype in a statistically enriched subset of genes that were DE by fasting (604 out of 925 genes, binomial test p-value < 2.2 x 10^-6^) (**Figure 4H**). Among those genes that were inversely regulated by fasting and *PER2-S662D*, *Clcn1* and *Ciart* were up-regulated by fasting, but down-regulated by *PER2-S662D* relative to wild type (**Figure 4I**). Conversely, *Cldn5* and *Wnt9a* were down-regulated by fasting, but up-regulated by *PER2-S662D* relative to wild type (**Figure 4I**). Importantly, RNA-sequencing in hypothalamus of *Per2* knockout mice revealed a direct relationship between *Per2* knockout and the genes that were DE by fasting for a statistically enriched subset of genes (714 out of 925 genes, binomial test p-value < 2.2 x 10^-6^) (**Supplemental Figure 4D and Supplemental Table 1**). For example, *Hmgcs2* is up-regulated both by fasting in wild type mice and by *Per2* knockout, while *Dbp* is down-regulated in both conditions (**Supplemental Figure 4E**). This result is consistent with our observations that *Per2* knockout decreased behavior during fasting while *PER2-S662D* increased behavior during fasting (**Figures 4D-E and Supplemental Figures 4B-C**). These results show that *PER2-S662D* opposes the hypothalamic transcriptional response to fasting that occurs in wild type mice at ZT16.

We next sought to understand which classes of fasting DE genes were blunted in expression in phospho-mimetic, *PER2-S662D* mice. We performed gene ontology analysis of the 604 genes that are DE by fasting in wild type mice and inversely regulated by the phospho-mimetic mutant (**blue fasting DE genes within Quadrants 2 and 4 of Figure 4H**). The top gene ontology terms that enriched from this analysis were generally classified into three categories: a) terms associated with kinase signaling cascades such as “EGFR tyrosine kinase inhibitor resistance” and “PI3K-AKT signaling pathway”; b) terms associated with polyunsaturated fatty acid metabolism such as “linoleic acid metabolism” and “alpha-Linolenic acid metabolism”; and c) terms associated more broadly with fatty acid metabolism such as “phospholipase D signaling pathway” and “ether lipid metabolism” (**Figure 4J**). Together, these data reveal a blunted transcriptional response to fasting in the hypothalamus of phospho-mimetic, *PER2-S662D* mutant mice, in-line with our behavioral and biochemical observations (**Figure 4K**). They suggest that fasting affects circadian behavior and hypothalamic transcription by hypo-phosphorylating PER2-S662.

### Polyunsaturated fatty acids regulate seasonal circadian entrainment

We identified a role for PER2-S662 phosphorylation in regulating gene expression programs associated with linoleic and linolenic acid metabolism during fasting. Linoleic acid (C18:2) and linolenic acid (C18:3) are the two most abundant polyunsaturated fatty acids (PUFAs) and serve as precursors for the synthesis of neuromodulatory and signaling compounds ^31^. PUFAs compete with monounsaturated fatty acids (MUFAs) for enzymatic binding, thus, the ratio of MUFAs to PUFAs, generally, and oleic acid (C18:1) to linoleic acid, specifically, regulates physiology and plays a role in metabolic diseases such as cirrhosis and diabetes ^31,32^. As linoleic and linolenic acids are essential fatty acids that must be taken up from the diet ^31^, we hypothesized that PUFA metabolism is a key nutrient-derived signal whereby the clock regulates circadian and seasonal behavior.

To first determine whether PUFAs and/or PUFA metabolites are modulated in the hypothalamus during fasting, we conducted unbiased lipidomics in hypothalamus and serum of *ad libitum* fed and fasted wild type mice at ZT14. We identified 618 lipid metabolites to be differentially abundant (“DA”, non-normalized differential analysis with DEseq2, FDR-adjusted p-value <0.05) in hypothalamus and 275 DA lipid metabolites in the serum (**Figure 5A, Supplemental Figure 5A, and Supplemental Table 2**). Remarkably, we detected numerous changes in PUFA metabolites in the hypothalamus during fasting, in-line with our transcriptomic findings (**Figure 4**). Specifically, the free fatty acid (FFA) and lysophosphatidylcholine (LPC) forms of linoleic and linolenic acids were among the most down-regulated DA metabolites by fasting in the hypothalamus, while the carnitine-conjugated forms (CAR) were among the most up-regulated in both compartments (**Figure 5A**). Free PUFAs in the hypothalamus were decreased by fasting during the dark period but not during the light period, consistent with effects from fasting and/or circadian timing (**Figure 5B**). Carnitine-conjugated PUFAs were elevated throughout the dark and light periods during fasting in the hypothalamus (**Figure 5B**). Triglycerides containing multiple unsaturated bonds were also decreased in the serum during fasting, but unaffected in the hypothalamus (**Supplemental Figure 5A and Supplemental Table 2**). Interestingly, we did not observe statistically significant changes in free linoleic or linolenic acids in the serum during fasting nor free oleic acid in either compartment (**Supplemental Table 2**). These results show alterations in PUFA metabolism in the hypothalamus and serum during fasting with notable depletion of PUFAs in the hypothalamus and accumulation of carnitine-conjugated PUFAs systemically.

**Figure 5:**
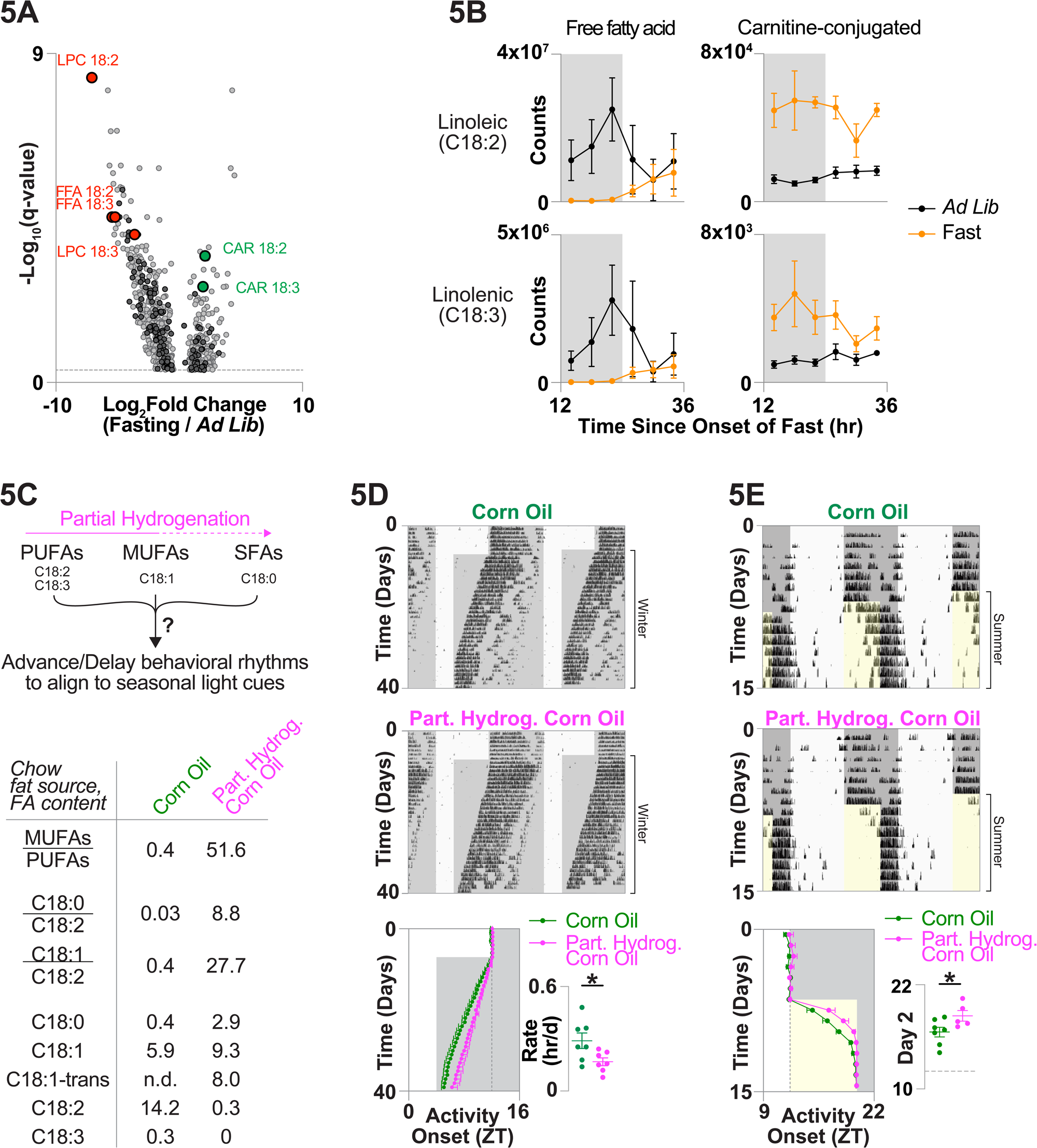
Polyunsaturated fatty acids regulate seasonal circadian entrainment. **A)** Lipidomics of hypothalamus from fed and fasted wild type mice at ZT14 showing differentially abundant metabolites (non-normalized DESeq2 FDR-adjusted p-value<0.05) that are unidentified (gray), identified (black), and of-interest (red and green). **B)** Quantification of free fatty acid (left) and carnitine-conjugated (right) forms of linoleic acid (top) and linolenic acid (bottom) across extended durations of fasting in hypothalamus of wild type mice. **C) (top)** Model of experimental design depicting use of isocaloric partially-hydrogenated fat chow to interrogate fatty acid saturation in seasonal circadian entrainment. **(bottom)** table of fat ratios and C18 fatty acid content (g/100g chow) in high-fat, isocaloric diets made from corn oil and partially-hydrogenated corn oil. **D and E)** Representative wheel-running traces and quantifications for **(D)** winter entrainment and **(E)** summer entrainment assays as described previously for mice given *ad libitum* access to chow containing corn oil or partially-hydrogenated corn oil. *p<0.05. Data are represented as mean +/-SEM. See also Supplemental Figure 5 and Supplemental Tables 2 and 3.

We were interested in interrogating the role of dietary PUFAs in nutrient-dependent entrainment to seasonal light cycles given their status as *essential* fatty acids. We had previously observed that HFD promotes entrainment to a summer light cycle and inhibits entrainment to a winter light cycle, while CR does the opposite (**Figure 1**). These phenotypes correlate with changes in calorie density and CK1δ activity, but they also correlate with the PUFA content of the chow and/or the organism. Relative to regular chow, HFD has increased content of numerous fatty acid species, including the C18 PUFAs: linoleic acid and linolenic acid (**Supplemental Table 3**). HFD also contains higher ratios of saturated fatty acids (SFA) to PUFAs, MUFAs to PUFAs, and oleic acid to linoleic acid than regular chow (**Supplemental Table 3**). During decreased calorie intake such as during chronic CR, PUFAs, MUFAs, and SFAs decrease in fat depots and the ratio of oleic acid to linoleic acid decreases in liver ^33,34^. Thus, we hypothesized that PUFAs, and/or the PUFA content relative to more saturated fatty acid species, regulate the rate at which mice entrain to seasonal light cycles.

To specifically interrogate the role of fat saturation in regulating entrainment to seasonal light cycles, we synthesized isocaloric high fat diets where the fat source was either corn oil (4.6 kcal/g, 45% calories from fat) or partially-hydrogenated corn oil (4.6 kcal/g, 45% calories from fat) (**Figure 5C**). Partial hydrogenation of corn oil maintains the same fatty acid content in terms of chain length, but hydrogenates fatty acids with multiple unsaturated bonds (PUFAs), converting e.g. C18:2 and C18:3 fatty acids into C18:0 and C18:1 fatty acids. Thus, the total PUFA content is lower in partially-hydrogenated corn oil relative to corn oil, while the MUFA/PUFA and oleic/linoleic ratios are higher. Specifically, high fat chow from corn oil has 14.2 grams of linoleic acid and 0.3 grams of linolenic acid per 100 grams of chow while high fat chow from partially-hydrogenated corn oil has 0.3 grams of linoleic acid and no detectable linolenic acid per 100 grams of chow (**Figure 5C and Supplemental Table 3**). Stearic acid (C18:0), oleic acid, and elaidic acid (C18:1-trans) are increased in partially-hydrogenated corn oil relative to corn oil. Thus, the MUFA/PUFA ratio is 51.6 in partially-hydrogenated corn oil chow and 0.4 in corn oil chow (**Figure 5C and Supplemental Table 3**). Similarly, the oleic/linoleic ratio is 27.7 in partially-hydrogenated corn oil and 0.4 in corn oil chow (**Figure 5C and Supplemental Table 3**). To test the role of fat saturation in seasonal circadian entrainment, we gave mice *ad libitum* access to these two diets and assessed the rate at which they entrained daily wheel-running behavior to light cycles that mimic winter and summer. Mice given the partially-hydrogenated corn oil diet with lower PUFAs and higher SFAs and MUFAs entrained to a winter light cycle at a slower rate (0.15 hours/day) than mice given the corn oil diet (0.25 hours/day) (**Figure 5D and Supplemental Figure 5B**). Indeed, 30 days after shifting lights to the 4:20 LD, mice given the partially-hydrogenated corn oil diet were 1.75 hours phase-delayed from mice given corn oil (**Supplemental Figure 5C**). Examining the effect of fat saturation on entrainment to a summer light cycle, partially-hydrogenated corn oil chow caused mice to entrain faster than mice given corn oil chow (**Figure 5E and Supplemental Figure 5D**). Indeed, two days after shifting lights to 20:4 LD, mice fed a partially-hydrogenated corn oil diet had a behavioral onset at ZT18.4 while mice fed a corn oil diet had a behavioral onset at ZT16.5 (**Figure 5E**). Together, these data show that an isocaloric diet with a partially-hydrogenated fat source promotes circadian entrainment to a summer light cycle and inhibits entrainment to a winter light cycle. Together with our findings with HFD and CR (**Figure 1**), these results suggest that the MUFA/PUFA ratio plays a key role in adapting circadian behavior to seasonal nutrient availability.

## DISCUSSION

### Nutrient abundance signals the changing of the seasons

Synchrony between circadian rhythms, food intake, and the external light/dark cycle is essential for maintaining organismal health ^35^, yet a gap remains in our understanding of the mechanisms whereby circadian rhythms adjust to the changing photoperiod and nutrient availability across the year. We found that nutrient availability alters circadian behavior to anticipate the season in which those nutrient conditions predominate (**Figure 6**). We utilized a high fat diet to mimic caloric-excess associated with long and intense daily light exposure during the summer, and calorie restriction to mimic nutrient limitation associated with short and weak daily light exposure during the winter. We found that these nutritional paradigms promote entrainment to the seasonal photoperiod in which they occur and inhibit circadian entrainment to the seasonal photoperiod in which they do not occur. It is intriguing to speculate that nutritional regulation of seasonal entrainment enhances evolutionary fitness by counteracting photic cues during seasons with atypical nutrient availability. In modern society where food availability for humans is constant throughout the seasons, one exciting area of current research involves the metabolic benefits of food consumption in specific phases of the day. These, “time-restricted feeding” and “intermittent fasting” interventions show promise in improving lifespan in mice ^36^ and metabolic health in mice^37,38^ and humans ^39,40^. Future research may interrogate the interplay between feeding phase, nutritional content, and the circadian clock in promoting healthy cognitive, metabolic, and physiological states across the seasons.

**Figure 6:**
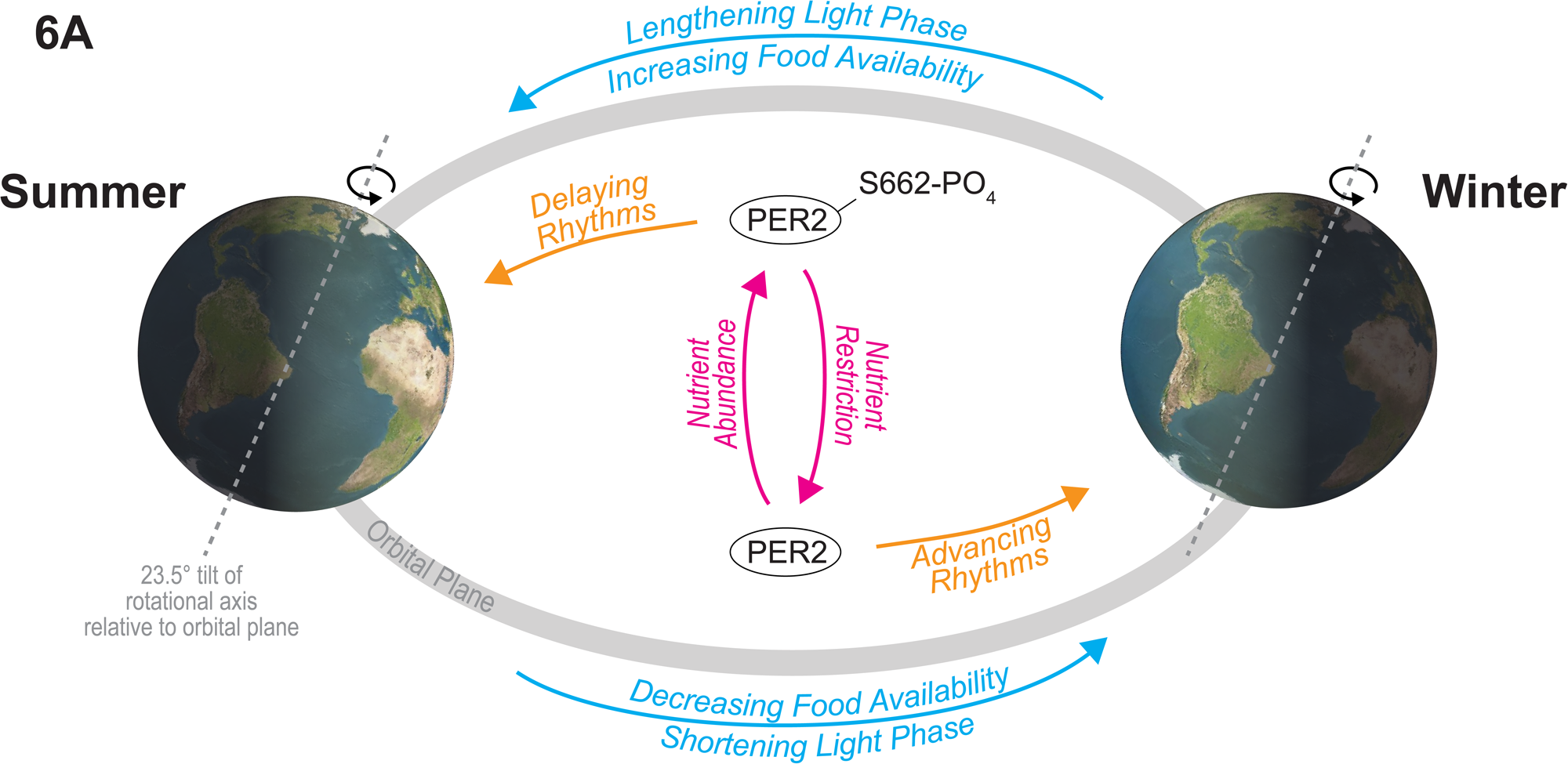
Model. **A)** The axial tilt of the Earth relative to the solar plane drives seasonal differences in nutrient availability and the duration of the daily light and dark phases. Animals must advance or delay circadian rhythms throughout the year to maintain synchrony between organismal rhythms and the light cycle. Nutrient content regulates PER2-S662 phosphorylation to control circadian phase and anticipate seasonal light cycles. Unseasonable availability of nutrient inhibits the behavioral phase-shifts that are required to maintain synchrony with the external light cycle.

### Interplay between circadian period and phase in nutritional seasonal entrainment

We have demonstrated that nutrient-density and content, PER2 dosage, and PER2-S662 phosphorylation regulate the rate of entrainment to seasonal photoperiods. We noticed that the genetic and nutritional conditions that drive phase advance and period shortening also promote entrainment to a winter light cycle, while those that drive phase delay and period lengthening promote entrainment to a summer photoperiod. HFD delays daily activity and feeding behavior in 12:12 LD cycles and lengthens the circadian period in constant darkness ^1^. *Per2* knockout shortens circadian period in mice housed in constant darkness before causing arrhythmia ^20^. *PER2-S662G* shortens circadian period in constant darkness and advances phase in 12:12 LD ^11^. Future efforts may ascertain the extent to which this observation can be generalized across other genetic and environmental interventions that affect circadian period or phase.

### PER2 is a circadian ‘monitoring station’ that integrates nutrient challenge

Our research has identified phosphorylation of PER2 on serine 662 as a key event that regulates entrainment of daily behavioral rhythms to seasonal light cycles during nutrient challenge. We previously identified this phosphorylation site from genetic interrogation of humans with FASP whose behavioral phase was advanced relative to the prevailing light cycle ^11,12^. PER2 is extensively post-translationally modified by phosphorylation ^16^. The PER2-S662 site, in particular, falls within a five-serine casein kinase target motif and within at least two competitive phosphoswitches ^15,41^. PER2-S662 phosphorylation is also regulated by acetylation/deacetylation of a nearby lysine (K680 in mice) that is in-phase with the casein kinase cluster ^13^. Finally, PER2-S662 is alternatively modified by O-linked GlcNAcylation, which prevents propagation of phosphoryl modifications on the five-serine cluster ^14^. The confluence of mass-action post-translational modifications from the metabolic state (acetylation, O-GlcNAcylation) with high-impact modifications (phosphorylation) and signal amplification mechanisms (multi-serine phosphorylation clusters and phosphoswitches), suggest that PER2 may act as a ‘monitoring station’ whereby cyclic modifications from the endogenous timekeeper compete with modifications that emanate from the environment and thus influence circadian timing. Certainly, we have found that PER2-S662 phospho-mutant animals are resistant to behavioral advance/delay caused by nutritional challenge, suggesting that this phosphorylation event sits at the nexus of these processes.

### Central and peripheral contributions to seasonality

Genetic manipulation of PER2 dosage and PER2-S662 phosphorylation revealed a role for the mammalian molecular clock, and PER2-S662 phosphorylation specifically, in regulating entrainment to seasonal light cycles. These studies provide novel insights into the interplay between the clock and metabolism, which was first uncovered with the discovery that *Clock* mutant mice develop metabolic syndrome and obesity ^42^. While the genetic paradigms that we used show a role for PER2 phosphorylation when it is altered across the whole organism, future studies will interrogate the role of PER2 and PER2-S662 phosphorylation in integrating metabolic signals in specific tissues throughout the organism. In the periphery, the liver clock may be a valuable context for study, as it rapidly senses the nutrient state ^43–46^, plays a key role in fatty acid liberation during hypo-caloric conditions ^47^, and is transcriptionally reprogrammed during the winter ^5^. In addition, the pancreatic β-cell rhythmically regulates lipolysis by circadian insulin secretion ^48^. Centrally, hypothalamic nuclei that regulate circadian rhythms, diurnal behaviors, and metabolism ^22,23,49^ are obvious candidates. Within the SCN, it may be intriguing to interrogate distinct sub-populations as the NMS-and VIP-expressing neurons regulate neurotransmitter plasticity in response to seasonal light cues ^50^. Neuroanatomical studies in *Drosophila* have similarly revealed divergent roles for the distinct subsets of clock neurons in control of seasonal rhythms ^10,51,52^. An interesting observation was that overexpression of the glycogen synthase kinase (GSK3) homolog, *shaggy*, in *Drosophila* ‘evening’ neurons is sufficient to drive rhythmic behavior in flies during constant light and summer-like photoperiods ^10^. We previously found that GSK3β phosphorylates OGT in mammals and drives PER2-S662 O-GlcNAcylation ^14^, suggesting a putative GSK3β / PER2 signaling axis that may be operative in mammalian neurons that regulate seasonality.

### PUFAs are nutritional signals that regulate diurnal behavior

We found that PER2-S662 phosphorylation regulates gene expression programs for linoleic and linolenic acid metabolism in the hypothalamus during fasting. These fatty acids are considered *essential* as they cannot be synthesized by mammalian metabolic enzymes and must be taken up from the diet ^31^. We found that altering the ratio of MUFAs to PUFAs in rodent chow by partially hydrogenating the fat source regulates the rate at which mice entrain to seasonal light cycles. Future studies are required to interrogate the potential mechanisms whereby PUFAs and MUFAs signal nutrient state to regulate daily behavior. Linoleic and linolenic acids serve as precursors for potent neuroactive and immunomodulatory molecules such as eicosanoids, prostaglandins, and endocannabinoids that may play a role in seasonal adaptation ^31,53^. Another particularly intriguing avenue of investigation may stem from the finding that linoleic acid, linolenic acid, and downstream molecules are endogenous ligands for the peroxisome proliferator activator receptor (PPAR) family of transcription factors that themselves regulate fatty acid metabolism ^54–57^. PPAR transcription factors are responsible for a portion of the daily oscillatory transcriptome by binding directly to PER proteins ^58,59^. It is possible that PER2-S662 phosphorylation and PUFAs affect seasonal behavior by altering activity or composition of hypothalamic PER/PPAR complexes, which would in-turn influence expression of PUFA metabolic genes and production of downstream signaling molecules. Detailed experiments will be required to tease apart the complex relationships in this system.

## ACKNOWLEDGEMENTS

We thank all members of the Ptáček and Fu laboratories as well as J.C., M.M., C.B.P., C.P., and J.B. for helpful discussions. Metabolomics data were acquired and processed at the UC Davis West Coast Metabolomics Center. This research was conducted while Daniel C. Levine was a Glenn Foundation for Medical Research Postdoctoral Fellow. Research support was provided by the National Institute of Neurological Disorders and Stroke (R01NS117929 to L.J.P. and R35NS132160 and R01NS104782 to Y.H.F); the Sandler Program for Breakthrough Biomedical Research, which is partially funded by the Sandler Foundation (to Y.H.F); the Novo Nordisk Foundation Project Grant in Bioscience and Basic Biomedicine (NNF17OC0028702 to T.M.P.); The Lundbeck Foundation Danish American Research Exchange Fellowship (to R.H.R.); and the Danish Cardiovascular Academy PhD Fellowship (PhD221006-DCA to R.H.R.)

## AUTHOR CONTRIBUTIONS

The design, execution, analyses, and visualization of all experiments were done by D.C.L. with assistance from T.M. in mouse work, and R.H.R., Y.H.F., and T.M.P. in data interpretation. L.J.P. supervised the project. D.C.L., L.J.P., and Y.H.F. wrote the manuscript.

## DECLARATION OF INTERESTS

The authors declare no competing interests.

## METHODS

### Lead Contact

Further information and request for resources and reagents should be directed to Louis Ptacek

### Materials Availability

No new materials or reagents were generated in this study.

### Data and Code Availability

RNA-sequencing data has been analyzed as above and has been deposited in the GEO public database managed by NCBI under the accession number GSE254155.

### Mice

All animal procedures were in accordance with guidelines of the Institutional Animal Care and Use Committee, and all mice were housed at 23-25C in the Rock Hall vivarium at the University of California San Francisco. Mice were of background B6/J and were maintained under 12hr light:12hr dark (LD) cycles with *ad lib* access to chow and water unless otherwise indicated. Male mice between 4 and 6 months old were used in experiments. *PER2-TgWT* mice containing a C-terminal FLAG tag were a gift from Dr. Ying Xu (Nanjing University). *Per2* knockout (stock 003819), *PER2-S662G* (stock 013700), and *PER2-S662D* (stock 013505) mouse lines were purchased from Jackson Laboratories.

### Wheel-running

Mice were singly-housed in cages containing wheels and maintained in light-proof boxes containing computer-controlled LED light sources set to an illumination of 100 lux (Phenome Technologies). Mice were acclimated to wheel-cages in 12:12 LD cycles for 1 week before lights were shifted to either a 4:20 LD cycle that approximates winter or a 20:4 LD cycle that approximates summer. Wheel-running activity and daily activity onset were determined with Actimetrics software. The rate at which mice entrain to a winter light cycle was taken by fitting a linear curve between the day that the lights were shifted and day +20 for all mice except *PER2-S662G*, which were day +2. Representative wheel-running traces are normalized by percentile.

### Dietary manipulations

High fat diets were purchased from Research Diets. Experiments with HFD in Figures 1 and 3 were performed with *ad libitum* access to a standard 45% high fat diet (D12451) containing a combination of lard (87.7%) and linseed oil (12.3%) and controlled with PicoLabs chow 5053 containing 13% calories from fat. Experiments with high fat diets containing corn oil and partially-hydrogenated corn oil in Figure 5 were sourced from Research Diets, contain the same composition and calorie content as D12451, but with the fat source listed, and were provided *ad libitum*. Calorie restriction was performed as previously using computer-assisted feeder devices produced by Phenome technologies and diets sourced from Bio-Serv ^47^. 300mg pellet versions of calorie restricted (F05314) and control (F05312) chows were purchased from Bio-Serv. The concentration of all macronutrients except sugar are supplemented to 1.6x for calorie restricted chow relative to control chow so that 40% less chow can be supplied to experimental mice to achieve a 40% calorie restriction. Mice were acclimated to feeders for 1 week with *ad libitum* access to control chow during 12:12 LD light cycles. For all dietary manipulations, regular chow was changed to the experimental chow ∼2 hours before the lights were shifted to the experimental condition. Amount and timing of calorie restricted and control chows were controlled by Actimetrics software and administered as described in the results.

### Video-based analysis of fasting behavior

Video-based activity was monitored as previously^60^. Briefly, mice were singly-housed and mice were allowed to acclimate to the new housing for 1 day. Activity was monitored over 6 subsequent days by infrared-cameras under constant infrared illumination. At ZT0 on the 5^th^ day, food, but not water, was removed from cages and bedding was changed before the cage was placed back in the same location in front of the IR camera. After 48 hours, food was replaced. Any-maze software (Stoelting Co) was used to track and report the minutes where the mouse was mobile within each 15-minute increment.

### RNA-sequencing and analysis

RNA-sequencing was performed from hypothalamic RNA extracts by Novogene. Briefly, RNA was extracted with Zymo directZol miniprep columns and resuspended in water. RNA meeting standard quality was prepared into libraries and sequenced on the Illumina NovaSeq to generate ∼20 million reads, 150 base pair, paired-end reads. STAR aligner was used to align fastq files to the mm10 genome, reads were assigned to genes with Subread::Featurecounts, and differential gene expression was determined with DESeq2 with an FDR-adjusted p-value cutoff less than 0.05 as previously ^13^.

### Immunoprecipitation & Western blotting

Immunoprecipitation and Western blotting were conducted as previously ^13^. Briefly, hypothalamus from *PER2-TgWT* mice that were fasted for 16 hours or given *ad libitum* access to HFD for 1 week was excised at ZT16 and extracted with ∼5 volumes of strong RIPA buffer containing kinase and phosphatase inhibitors (Abbexa abx090624), sonicated in a water bath 3 x 30sec on high, quantified by DC protein quantification kit (Bio-Rad) and diluted to 4 mg/mL in RIPA. 100 µL of lysate was diluted with 900 µL of dilution buffer (1.1% triton X-100, .01% SDS, 1.2 mM EDTA, 16.7 mM Tris, 167 mM NaCl), and 25 µL of BSA-blocked anti-FLAG conjugated magnetic beads (Sigma) were incubated overnight at 4C, rotating end over end. Immunoprecipitates were eluted by boiling, then loaded into 5% SDS-PAGE gels. Western blots were performed by transferring gels to nitrocellulose membranes and blocking in 5% BSA in PBST. Rabbit anti-PER2 and rabbit anti-PER2-S662p antibodies developed previously ^12,14^ were used at 1:2000 and 1:500, respectively. Anti-rabbit secondary antibodies were used at 1:2000 and membranes exposed to film and developed.

### Lipidomics

Hypothalamus and serum samples as described in the text were submitted for complex lipids analysis at West Coast Metabolomics Center. Samples were extracted using Matyash extraction procedure, which includes MTBE, MeOH, and H2O. The organic (upper) phase was dried down, then resuspended in 110 uL of a solution of 9:1 methanol: toluene and 50 ng/mL CUDA. Following centrifugation, each sample was aliquoted into three parts. 33 uL are aliquoted into a vial with a 50 uL glass insert for positive and negative mode lipidomics. The last part is aliquoted into an eppendorf tube to be used as a pool. The samples are then loaded in an Agilent 1290 Infinity LC stack. The positive mode was run on an Agilent 6530 with a scan range of m/z <u>120-1200</u> Da and an acquisition speed of 2 spectra/s. Positive mode has 1 uL injected onto an Acquity Premier BEH C18 1.7 µm, 2.1 x 50 mm Column. The gradient used is 0 min 15% (B), 0.75 min 30% (B), 0.98 min 48% (B), 4.00 min 82% (B), <u>4.13-4.50</u> min 99% (B), <u>4.58-5.50</u> min 15% (B) with a flow rate of 0.8 mL/min. The other sample aliquot was run in negative mode, which was run on Agilent 1290 Infinity LC stack, and injected on the same column, with the same gradient and using an Agilent 6546 QTOF mass spec. The acquisition rate was 2 spectra/s with a scan range of m/z 60-1200 Da. The mass resolution for the Agilent 6530 is 10,000 for ESI (+) and 30,000 for ESI (-) for the Agilent 6546. Differential metabolites were determined with DESeq2 with normalization factors set to 1 and with an FDR-adjusted p-value cutoff < 0.05.

### Statistics and data representation

Data are represented by mean +/-standard error of the mean and statistics were performed by two-tailed, unpaired student’s t test except where otherwise noted in results, figure legends, or methods. Differences were considered significant when p < 0.05 except where otherwise noted in results, figure legends, or methods.

## Notes

https://www.ncbi.nlm.nih.gov/geo/query/acc.cgi?acc=GSE254155

### Competing Interest Statement

The authors have declared no competing interest.

