## Supplementary figures and images for "Nutrient Abundance Signals the Changing of the Seasons by Phosphorylating PER2"

### Supplemental Figures

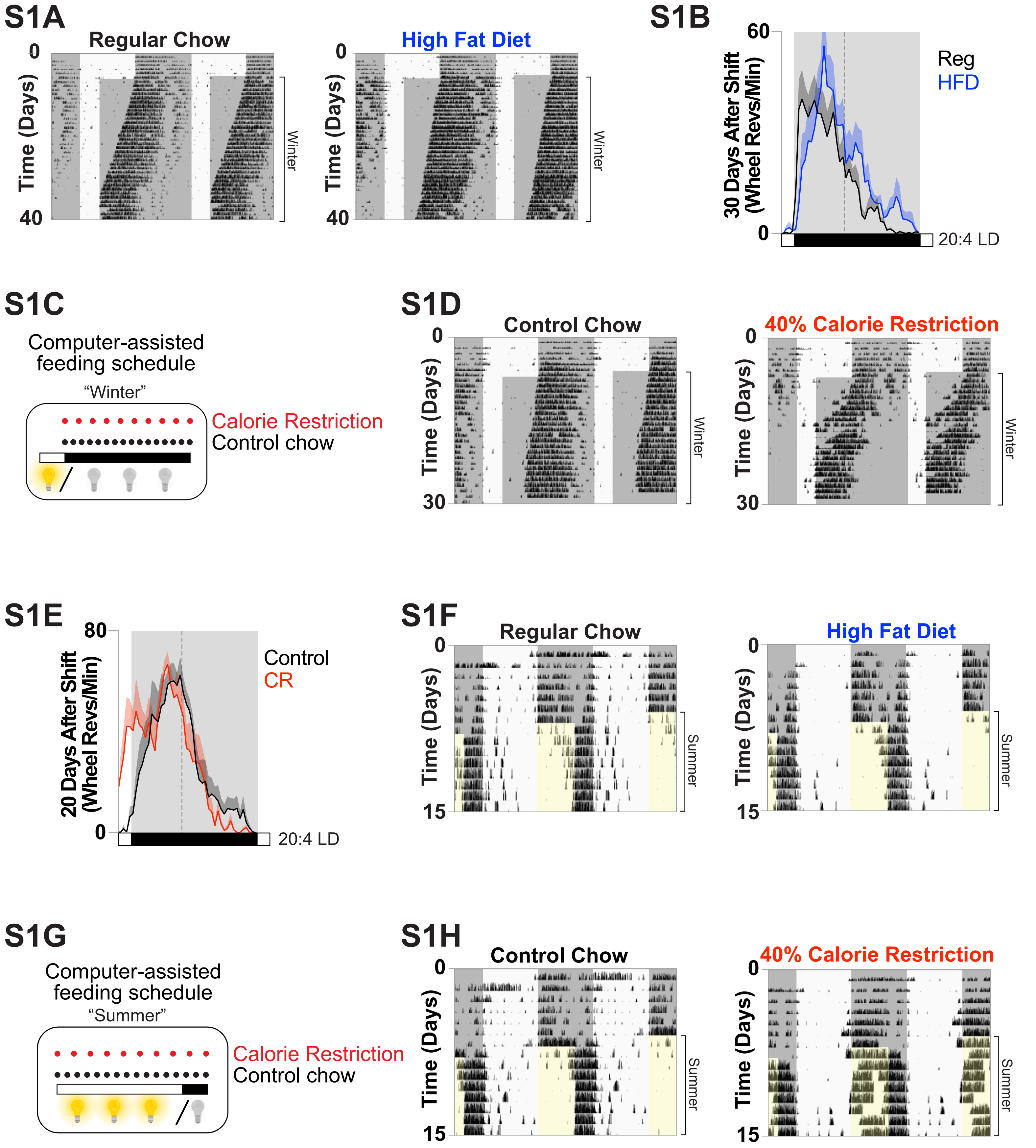

## S2A

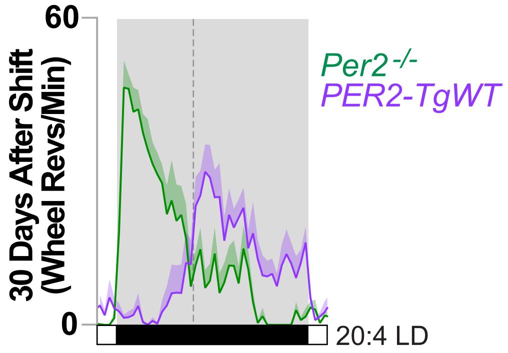

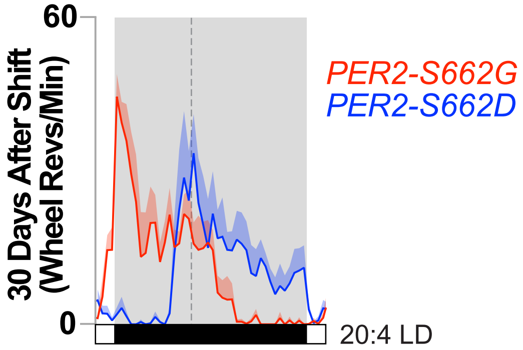

S4A

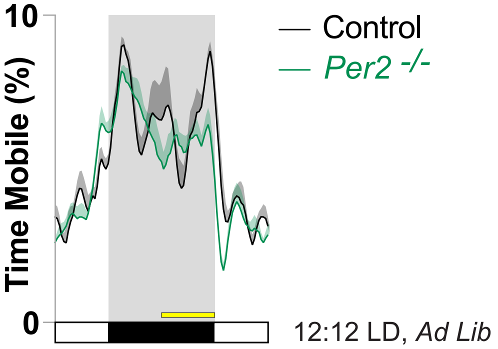


S4B

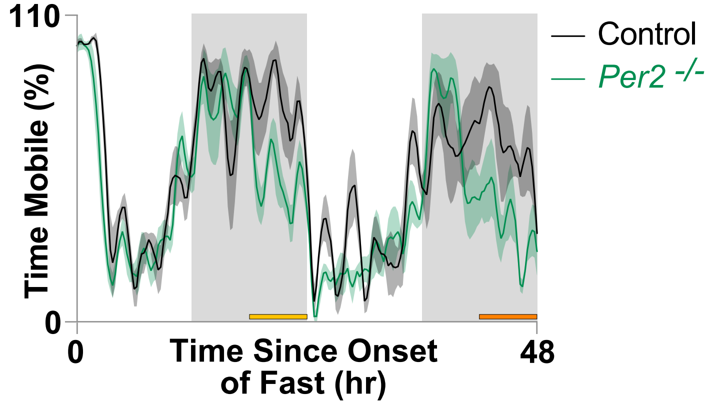


S4C

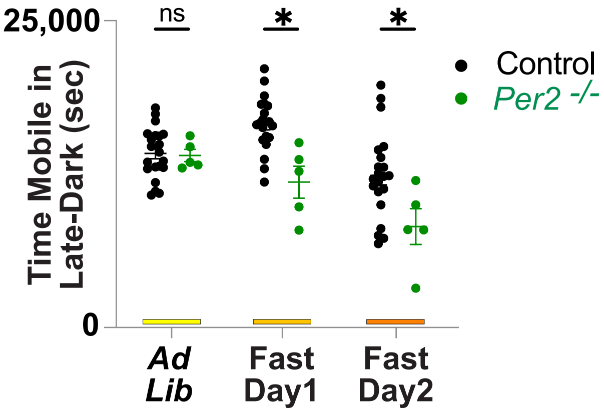


S4D

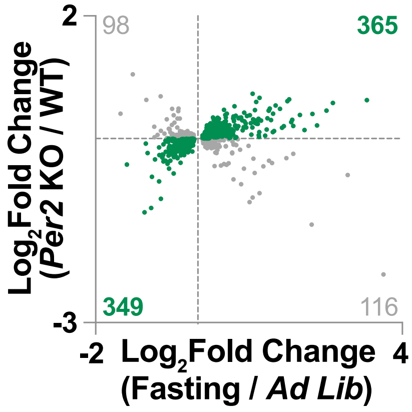


S4E

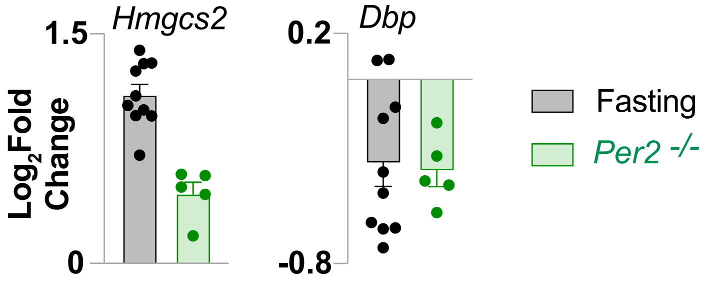

S5A

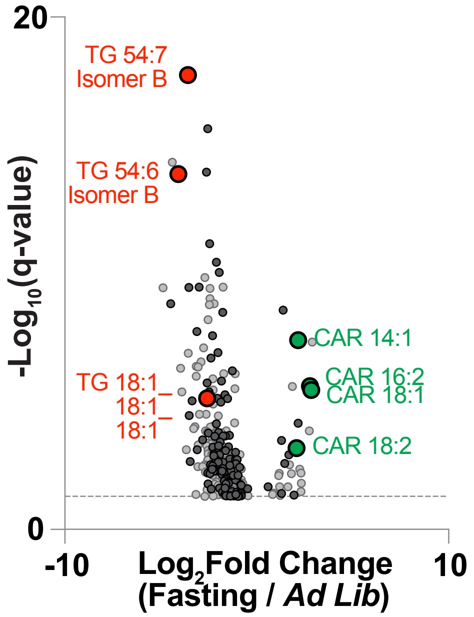


S5D

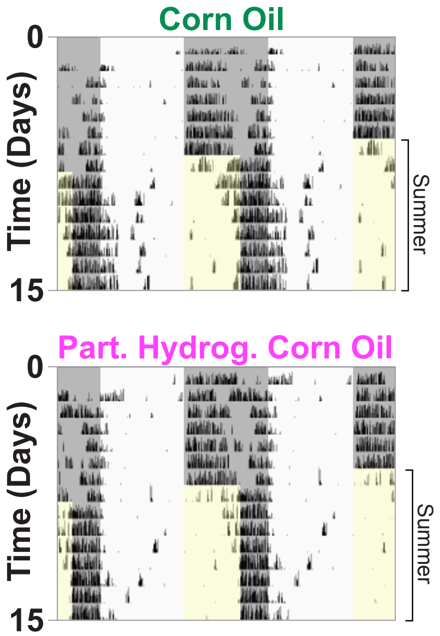


S5B

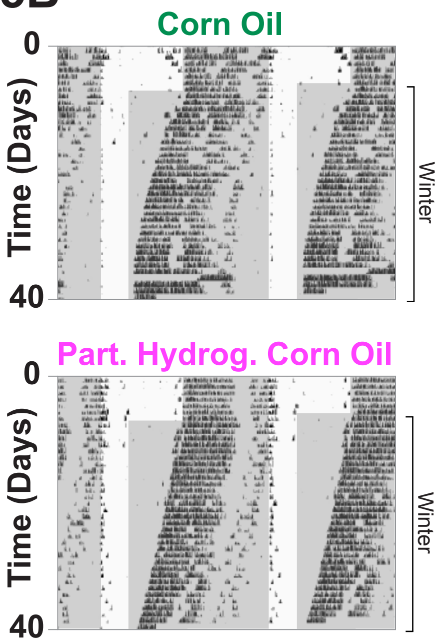


S5C

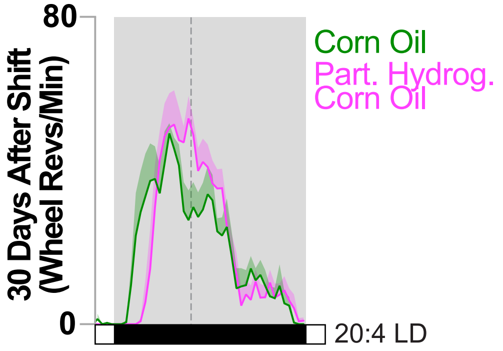
